# Proteomic analysis of the TDP-43-associated insoluble fraction from NEFH-TDP-43 mouse brain suggests sustained stress granule formation, CLUH granule recruitment and impaired mitochondrial metabolism

**DOI:** 10.1101/2024.05.23.595607

**Authors:** Felicity Dunlop, Shaun Mason, Stavroula Tsitkanou, Aaron P. Russell

## Abstract

Cytoplasmic accumulation and aggregation of TDP-43 is a hallmark of ∼97% of ALS cases. Formation of TDP-43 insoluble aggregates is suggested to either directly or indirectly cause motor neuron loss and progressive neuromuscular degeneration, although how this occurs is not precisely understood. Cytoplasmic TDP-43 is observed in stress granules (SG). SGs are ribonucleoprotein (RNP) complexes formed during stress conditions, consisting of mRNAs and RNA-binding proteins (RNPs). Chronic TDP-43/SG formation may play a role in neuromuscular degeneration in ALS. The composition of *in vivo* TDP-43-asscociated SGs in ALS not known. This knowledge may provide insights into the molecular pathways impaired by TDP-43-associated SGs and suggest disease modifying mechanisms. The aim of this study was to isolate and analyse the proteome of the insoluble TDP-43-associated SG fraction from brain tissue of end-stage TDP-43ΔNLS mice. Proteomic analysis identified 134 enriched and 17 depleted proteins in the TDP-43ΔNLS mice, when compared to the control mice. Bioinformatics analyses of the impacted proteins from the SG preparation suggested that brain tissue from end-stage NEFH-TDP-43ΔNLS mice have sustained SG formation, CLUH granule recruitment and impaired mitochondrial metabolism. This is the first time that CLUH granule recruitment has been demonstrated in ALS and the known role of CLUH suggests that cell starvation is a potential mechanism of motor neuron loss that could be targeted in ALS.

**Highlights:**

1. We present a detailed a protocol for the extraction of cross-linked TDP containing stress granules from brain tissue.
2. We present proteomics data from the insoluble fraction from brain tissue of an ALS mouse model.
3. We identify the mitochondrial mRNA transport protein CLUH and CLUH targets trapped in insoluble SG fraction of brain.
4. Reanalysing proteomics data from axonal soluble fraction supports a link between proteins trapped in the brain and depleted from the axons.
5. Propose a model where metabolic mitochondrial enzymes trapped in the insoluble fraction from the brain via a CLUH dependent mechanism results in motor neuron death by starvation in ALS.

## Introduction

The NEFH-TDP-43ΔNLS mouse is a well-established model of amyotrophic lateral sclerosis (ALS). Disruption of the nuclear localisation sequence of the native human TDP-43 coding sequence results in cytoplasmic accumulation of TDP-43 in neuronal tissues (Walker et al., 2015), which is a hallmark of ∼97% of ALS cases (Ling et al., 2013; Neumann et al., 2006). Cytoplasmic TDP-43 forms clumps of protein within the cell which characteristically pellet with centrifugation unless extracted by boiling in SDS first (Xiong et al., 2014). The insoluble aggregate results either directly or indirectly in motor neuron loss and progressive neuromuscular degeneration (Arai et al., 2006; Hergesheimer et al., 2019; Neumann et al., 2006). The intracellular aggregation of insoluble TDP-43 contributes to its broad and complex role in ALS as it impacts numerous processes within multiple compartments, including the nucleus, cytoplasm, mitochondria and neuromuscular junction (Lepine et al., 2022; Prasad et al., 2019; Wang et al., 2019). TDP-43 protein aggregation is theorised to occur via interaction of the intrinsically disordered c-terminal domain (Prasad et al., 2019; Santamaria et al., 2017) or via the phosphorylation of TDP-43 (Hasegawa et al., 2008; Inukai et al., 2008; Neumann et al., 2006); the latter influencing liquid-liquid phase separation and interaction with intracellular stress-related entities including stress granules (SG) (Colombrita et al., 2009).

SGs are membraneless ribonucleoprotein (RNP) complexes formed by the aggregation of mRNAs and RNA-binding proteins (RBPs) (Advani & Ivanov, 2019). SGs are formed during stress conditions and increase in size and number in response to stress, including heat shock, oxidative stress, mitochondrial stress (Kedersha et al., 2005; Ohn et al., 2008). Canonical SG formation is initiated by phosphorylation of eukaryotic initiation factor 2α (eif2α) by the 4 known stress specific kinases PERK (unfolded proteins), HRI (oxidative stress), GCN2 (starvation) or PKR (viral infection/cytoplasmic RNA) (Matsumiya et al., 2023). The physiological role of SGs is to protect cytoplasmic untranslated mRNAs and stalled pre-initiation complexes following acute stress by protein-protein and protein-RNA binding (Wolozin & Ivanov, 2019). These acute granules dissipate following stress removal. However, their chronic presence results in pathological cytoplasmic aggregates, that have been linked to ALS (Daigle et al., 2016). TDP-43 is a major architect in the development of physiological SGs and in pathological cytoplasmic aggregates in neurons of people with ALS and in other neurodegenerative conditions ALS (Fan & Leung, 2016; Prasad et al., 2019). TDP-43 contributes to SG development in neuronal tissue through its assembly with core RBPs such as T-Cell-Restricted Intracellular Antigen- (TIA-1), eukaryotic translation initiation factor (eIF3), polyadenylate-binding protein (PABP) and GTPase Activating Protein (SH3 Domain) Binding Protein (G3BP1) and 40S ribosomal subunits (Fan & Leung, 2016; Wolozin & Ivanov, 2019) and through the formation of amyloid-like aggregates (Prasad et al., 2019).

Our current understanding of formation, composition and potential role of SGs in ALS has come from cell culture studies (Asadi et al., 2021). Several studies have also performed histological analysis of post-mortem brain samples from ALS donors, identifying TDP-43 cytoplasmic inclusions colocalizing with SG markers, eIF3, G3PB1, PABP-1 and TIA-1 (Asadi et al., 2021; Liu-Yesucevitz et al., 2010). Attempts to determine the presence of TDP-43 in SG fractions isolated from tissues is limited and may lack specificity due to TDP-43 immunoprecipitation from soluble and not insoluble fractions (Montalbano et al., 2020). Proteomics analysis has been performed to observe changes in postmortem frozen motor cortex samples form ALS donors compared with healthy control samples (Lee et al., 2023) as well as longitudinal analysis in cortex samples from NEFH-TDP-43 mice (San Gil et al., 2024). These studies identified multiple disease-related changes in biological pathways. However, these analyses were performed using soluble protein fractions (Lee et al., 2023; San Gil et al., 2024) without TDP-43 immunoprecipitation, and would therefore not accurately capture the TDP-43-associated SG proteome.

Presently, the composition of TDP-43-associated SGs from *in vivo* models of ALS is lacking. Here, we prepared and analysed the proteome of the insoluble TDP-43-associated SG fraction from brain tissue of end-stage TDP-43ΔNLS mice. When compared with) control mice, 134 proteins were enriched and 17 were depleted in the SG fraction. Bioinformatics analyses of the impacted proteins suggested the brain tissue from end-stage TDP-43ΔNLS mice have sustained SG formation, CLUH granule recruitment and impaired mitochondrial metabolism.

## Methods

### Animals

Six hTDP-43ΔNLS transgenic (Tg) mice harbouring a doxycycline (DOX)-suppressible transgene to express human TDP-43 (hTDP-43) with a defective nuclear localization signal (hTDP-43ΔNLS) under the control of the human neurofilament heavy chain *NEFH* promoter and 7 age-matched control mice were used in this study. The NEFH-TDP-43 mice were produced by the inter-cross of C57/BL6 *NEFH*-tTA^+/−^ mice and C57/BL6 *tetO*-hTDP-43ΔNLS^+/+^ mice (Tsitkanou et al., 2022; Walker et al., 2015). All mice were maintained on DOX-enriched chow (200 mg/kg; Gordons Specialty Feeds, Bargo NSW) and genotyped when weaned (Walker et al., 2015). DOX-enriched chow was removed at 6 weeks of age to induce the transgene. Seven littermate single transgenic mice were used as healthy, control mice and remain unaffected with the removal of the DOX-enriched chow. Mice were humanely killed 5 weeks after disease initiation by CO_2_ asphyxiation when they approach disease end-stage and were close to 20% body weight loss. Mice were maintained on 12 hours light/dark cycles at a temperature range of 18-24°C and humidity between 40-70%. Brains were immediately removed and snap frozen in liquid nitrogen and stored at −80°C until required. The study was approved by the Deakin University Animal Ethics Committee (G21-2020).

### Immunoprecipitation and Western Blotting

For the initial antibody titration experiment, RIPA brain tissue extracts were prepared as previously described (Tsitkanou et al., 2022). Each IP (immunoprecipitation) contained 100 µg of RIPA extracted protein that was precleared with protein G magnetic beads.

Preclearing lysates with Protein G magnetic beads: Protein G magnetic beads (10 µl per sample) were washed with 300 µl modified RIPA buffer (50 mM Tris-HCl, pH 7.4, 150 mM NaCl, 0.25% deoxycholic acid, 1% NP-40, 1 mM EDTA, Millpore cat# 20-188) + 1:100 HALT Protease and Phosphatase inhibitors (cat# 78438, Pierce)) for 5 minutes on a rotating wheel at 4⁰C in individual 1.5 ml Eppendorf tubes. The washed beads were then placed on a magnetic rack (cat# 12321D, Invitrogen) to remove wash buffer. RIPA buffer and RIPA brain lysate (100 µg in a 300 µl final volume) were added to the beads. Following this 10 µl of 6% BSA was added and mixed on a rotating wheel at 4°C for 1-2 hours.

Immunoprecipitation: The precleared lysates were transferred to fresh 1.5 ml Eppendorf tubes containing 20 µl RIPA washed Protein G magnetic beads and 0.5, 1.0, or 2.0 µg of mouse IgG1 anti-human TDP-43 antibody (ProteinTech cat# 60019-2-Ig) was then added to each tube. The samples were placed on a rotating wheel overnight at 4°C. The following day samples were placed on magnetic rack for 2 minute to allow for bead binding. The supernatant was removed, and the beads washed a total of five times (Thomas et al., 1998). Beads were washed 2 x with wash buffer I (50 mM Tris-HCl, pH 7.5, 150 mM NaCl, 1% Triton X-100, 0.5% sodium deoxycholate), + inhibitors (cat# 78438, Pierce) 2 x with wash buffer II (50 mM Tris-HCl, pH 7.5, 500 mM NaCl, 0.1% Triton X-100, 0.05% sodium deoxycholate) and 1 x with wash buffer III (50 mM Tris-HCl, pH 7.5, 0.1% Triton X-100, 0.05% sodium deoxycholate). Samples for Western blotting were resuspended in 15 µl Urea-based SDS sample buffer (62.5 mM Tris-HCl, pH 6.8, 2% SDS, 10%-mercaptoethanol (vol/vol), 6 M urea, 20% glycerol), (Thomas et al., 1998) and heated at 95⁰C for 5 minutes. 30µg of RIPA lysate (starting material was included as a Western blot positive control) and samples were resolved on 4-15% Criterion Stain free gels (cat# 5678085 Biorad) at 200 Volts, 40 minutes. Proteins were transferred to a PVDF membrane by Western blotting at 100 volts for 1 hour. Membranes were blocked in 5% BSA in PBS at room temperature for 1 hour before overnight incubation with primary antibody mhTDP-43 (rabbit; Cat#10782-2-AP; 1:1000; ProteinTech). Blots were washed 3 times in TBST (20 mM TRIS, 150 mM NaCl, 0.1% w/v Tween^®^ 20 detergent) for 10 minutes each. Donkey anti-mouse AlexaFluor 800 IgG antibodies (Life Technologies) diluted at 1:10,000 in 5% BSA in PBS was used as the secondary antibody with incubation at room temperature for 1 hour. Blots were washed twice in TBST and once in PBS and then imaged using the infrared imaging system (Li-COR Biosciences, Lincoln, NE, USA).

### Formaldehyde crosslinking and tissue lysis

Formaldehyde crosslinking and homogenization of brain tissue was adapted from previous protocols (Liu-Yesucevitz et al., 2010; San Gil et al., 2024; Sap et al., 2021). 500 µl of 0.5% formaldehyde in PBS was added to 30 mg of frozen mouse brain tissue, previously crushed under N_2_ and placed on a rotating wheel for 10 mins at room temperature. Samples were then centrifuged at 6000 g for 15 seconds followed by removal of formaldehyde. To quench crosslinking, 500 µl of 250mM Tris-HCl pH 7.5 was added to the sample and gently mixed by pipetting for 45 seconds using wide bore pipette tips. The samples were then pelleted at 6000 g for 15 seconds followed by removal of Tris. 500 µl of ice-cold Co-IP buffer (20 mM Tris-HCL pH 7.5, 150 mM NaCl, 1 mM EDTA, 1% Triton X-100 + protease and phosphatase inhibitors) was then added to the pellet and placed on ice. Five 1.4mm Precellys beads (Bertin Instruments, Montigny-le-Bretonneux France) were added to each tube followed by homogenisation (Precellys bead homogeniser + CryoLys; Bertin Instruments, Montigny-le-Bretonneux, France) at 6000 rpm for 30 seconds each, with liquid N_2_ cooling to 4⁰C. Following this, the samples were de-foamed in a CentriVap (Labconco, Kansas, USA) for 15 seconds. The homogenisation and de-foaming steps were repeated 5 times. The samples were then placed on rotating wheel for 1 hour at 4⁰C.

### Stress Granule Enrichment

Stress granule enrichment was based on previous protocols (Jain et al., 2016; Wheeler et al., 2017). Following lysis the samples were centrifuged at 1000g for 5 min at 4⁰C with the supernatant then decanted into a 1.5 ml Eppendorf tube and the pellet discarded as cellular debris. The supernatant was then centrifuged at 18,000 g for 20 min at 4⁰C. The stress granule (SG) pellet was resuspended in 500 µl Co-IP buffer and centrifuged again at 18,000 g 20 min at 4⁰C. The SG pellet was again resuspended in 500 µl Co-IP buffer. A 5 µl aliquot was removed and used to determine protein concentration in triplicate using the BCA assay (Cat# 23225, Pierce). The remaining SG sample was kept on wet ice until required for immunoprecipitation.

### TIA-1 and TDP-43 Immunoprecipitation of Crosslinked Samples

Bead washing and preclearing was conducted the same as for the non-cross-linked (native) samples above, with 10 µl of protein G magnetic beads per sample. Co-IP buffer was used in place of RIPA buffer and 150 µg stress granule lysate was used. Immunoprecipitation was conducted the same as for native samples with 2.0 µg of either mouse IgG1 anti-human TDP-43 antibody (ProteinTech cat# 60019-2-Ig) or TIA-1 (Santa Cruz cat#1751) used for each overnight immunoprecipitation. Following the immunoprecipitation samples were washed using high stringency “SAP” wash buffers adapted from (Sap et al., 2021). Beads were washed 2 x with wash buffer I + inhibitors (50mM Tris pH 7.4; 300mM NaCl; 1mM EDTA; 0.1% Triton X-100), 1 x with wash buffer I (no inhibitors) and 1 x with wash buffer II (50mM Tris pH 7.4; 150mM NaCl; 1mM EDTA; 0.01% Triton X-100). Western blotting of crosslinked samples was conducted the same as for native samples but with samples heated at 100⁰C for 20 minutes in urea-based SDS sample buffer to reverse the crosslinking prior to SDS-PAGE.

### Sample Preparation for Proteomics

SAP washed beads of immunoprecipitated samples for proteomics analysis were resuspended in 90 µl 2% SDS, 62.5 mM Tris-HCl (pH 6.8)(Sap et al., 2021). Following this 1.8 µl of 0.5 M TCEP (cat #C4706) and 8 µl 0.5 M IAA (cat #I6125) were added separately to the resuspended beads. Single step formaldehyde crosslinking reversal and disulfide bond reduction and alkylation was then done by heating the samples at 100⁰C for 30 minutes in the dark. The samples were cooled on ice and then placed on the magnetic rack for 2 minutes before the eluted proteins were transferred to a fresh 1.5ml tube and the spent beads discarded.

SP3 and Trypsin digestion: The eluted proteins were prepared for trypsin digestion using SP3 single-pot, solid phase enhanced protocol (Hughes et al., 2019). 5 mg of each Sera-Mag Hydrophilic and Hydrophobic magnetic SpeedBeads (Cytiva, cat. no. 45152105050250, cat. no. 65152105050250) were combined and washed with 1 ml LC-MS water (cat #1.15333) on a magnetic rack. Washed beads were resuspended in 1 ml LC-MS water and 20 µl added to each tube of eluted proteins. Binding of the proteins to the beads was induced with 120 µl of absolute ethanol (cat #459836) and mixed using a Thermomixer (Eppendorf, Macquarie Park, Australia) at 1600 rpm for 8 minutes. The bead bound proteins were washed three times with 80 % ethanol before resuspension in 60 µl digestion buffer (10% TFE, 100 mM HEPES pH 7.3) and 0.1 µg trypsin (cat #) and incubated for 18 hours at 37⁰C, 1200 rpm using a Thermomixer.

STAGE tips: Overnight trypsin digests were desalted using STAGE tips prepared from 2 layers of SDP-RPS membrane (Cat # 2341 Empore). The STAGE tips were first equilibrated in 50 µl each of acetonitrile (Cat #1.00029), 0.2 % TFA (Cat# T0699) in 30% methanol (cat #1.06035), and then 0.2% TFA by centrifugation at 500 g for 3 minutes each. Overnight trypsin digests were placed on the magnetic rack. The supernatant containing the digested peptides were pipetted on top of equilibrated STAGE tip containing 150 µl 1 % TFA and centrifuged at 1000 g for 8 minutes. Tips were washed twice with 100 µl 0.2% TFA and once with 40 µl 1% TFA in 90% isopropanol by centrifugation at 1000 g for 5 minutes. Peptides were eluted into a fresh 1.5 ml tube with 60 µl 5 % ammonium hydroxide (cat # 221228) in 80 % acetonitrile and centrifuged at 1000 g for 3 minutes. Eluted peptides were dried in a CentriVap (Labconco, Kansas, USA) at 45⁰C for 30 minutes until dry and stored at 4⁰C. Peptides were resuspended in 20 µl loading buffer (2% (v/v) acetonitrile, 0.1% (v/v) TFA) for mass spectrometry analysis.

### Liquid chromatography mass spectrometry

Samples were analyzed using an UltiMate 3000 RSLCnano (ThermoFisher Scientific) linked to an Orbitrap Exploris™ 240 Mass Spectrometer (ThermoFisher Scientific). Six microlitres of sample was loaded onto a PepMap 100 C18 (20 mm x 0.1mm, 5µm) nanoViper trap column for 3 mins at a flow rate of 15 µL/min with 2% (v/v) acetonitrile, 0.1% (v/v) TFA, and then resolved on a PepMap Neo C18 (75 µm x 500 mm, 2µm) nanoViper analytical column at a flow rate of 250 nL/min. Mobile phases consisted of 0.1% (v/v) Formic acid (A) and 80% (v/v) Acetonitrile/0.1% (v/v) Formic acid (B). Gradient conditions were: 0-2 min 2.5% B, 2-2.1 min 8% B, 2.1-122 min 32% B, 122-125 min 45% B, 125-133 min 99% B, 133-150 min 2.5% B. The column compartment was set at 50°C. Ion source conditions were: NSI type, static spray voltage, 2000V positive ion, 1500V negative ion, 275°C ion transfer tube temperature. MS1 data were acquired over 350-1200 m/z with an orbitrap resolution of 120000, RF lens 70%, and positive polarity. Dynamic exclusion was for 20s, with mass tolerances of 10 ppm. Data dependent analysis used a 3 s cycle time with HCD collision energy set at 30%. MS2 spectra were acquired with a fixed first m/z of 120 with an orbitrap resolution of 15 000.

### Data analysis

Data were processed using Thermo Proteome Discover (v.3.0) with Sequest HT as the database search algorithm against the *Mus musculus* FASTA file (Uniprot). Input data to Sequest HT were as follows: Maximum missed tryptic cleavages 2, precursor mass tolerance 10 ppm, fragment mass tolerance 0.02 Da. Dynamic modifications were oxidation (methionine), and static modifications were carbamidomethylation (cysteine). The processing workflow also had the Percolator node used, with strict target FDR set at 0.01 and relaxed FDR set at 0.05. The target/decoy strategy was concatenated with validation based on the q-value. Data Normalization based on total peptide amount was used to compensate for sample-to-sample variation. Protein abundances were calculated on the basis of summed peptide abundances of unique and razor peptides.

A complete list of all the TDP-43 associated proteins was generated by selection of all proteins that had quantifiable protein abundances in the TDP-43 sample group based on the unique accession numbers of the master protein. The gene symbols were used for cross species comparison against the Lunenfield RNA granule database and specific stress granule lists from (Jain et al., 2016) and (Hu et al., 2023). Protein ratios were calculated using protein abundances and Hypothesis testing was based on ANOVA (individual protein) calculations. Fold change individual protein ratios TDP-43/WT of >1.5 (Log2=0.58) and ANOVA p-value <0.05 were used as statistical cutoffs to determine significance and no imputation was used.

Protein-protein interactions, GO terms and Reactome pathways were identified using STRING (https://string-db.org/). STRING was used for analysis as it is a freely available resource, allowing other researchers to reproduce the analysis from the list of proteins in Supplemental Tables 2 and 3. A CLUH target list was generated by combining the data from (Gao et al., 2014) and (Schatton et al., 2017). All 256 proteins were retrieved from Table S3 (Guo et al., 2017) and 167 proteins were retrieved by sorting Table S1 data for Log2 Ratio KO/WT >0.58 and p-value <0.05 from Schatton *et al* (Schatton et al., 2017). The lists were combined as gene symbols to allow for comparison with both mouse and human proteins. Duplicates removed to provide a final list of 371 unique CLUH target proteins.

Supplemental Table S6 (file mmc6) was retrieved from (Guise et al., 2024) and the results sorted for p-value <0.05, Log2(FC)<0.58 for ALSvsCTL. The dataset described as “decreased early and late” was set to “true” to retrieve 540 consistently depleted proteins. The 41467_2021_27221_MOESM5_ESM file from (Altman et al., 2021) containing 249 depleted proteins was accessed and the gene symbol and corresponding accession numbers for the Log2FC<-1 246 proteins were retrieved (3 entries were removed as they were incomplete). Gene symbols were mapped to unique accession numbers using the Uniprot ID mapping function (https://www.uniprot.org/id-mapping) to prevent ambiguity. Mitochondrial proteins from the Guise *et al* dataset were then identified by comparison with the human mitochondrial database (Human MitoCarta3.0: 1136 mitochondrial genes: https://personal.broadinstitute.org/scalvo/MitoCarta3.0/human.mitocarta3.0.html) using the unique accession numbers. The same identification method was used for the Altman *et al* dataset however comparison was made with the mouse mitochondrial database (Mouse MitoCarta3.0: 1140 mitochondrial genes: https://personal.broadinstitute.org/scalvo/MitoCarta3.0/mouse.mitocarta3.0.html). Both datasets were also checked against the CLUH target list. The CLUH target proteins from both datasets were then submitted to the STRING database as accession numbers for Reactome Analysis.

## Results

### Establishing the stress granule enrichment and TDP-43 immunoprecipitation technique

The specificity of our immunoprecipitation (IP) with the TDP-43 antibody recognizing the human isoform (hTDP-43) is shown in Figure 1A. A TDP-43 IP was performed on RIPA lysates from crushed brain tissue from the NEFH-TDP-43 and Control mice. This was performed using increasing amounts of the hTDP-43 antibody (0.5, 1.0 and 2.0 µg). The IP samples were then probed using a TDP-43 antibody recognizing both the mouse and the human isoforms (mhTDP-43). RIPA extracted lysates (30 µg starting material) from the NEFH-TDP-43 and control mice were used as positive control samples. A band was detected in the RIPA extracted lysate from the NEFH-TDP-43 and control mice when using the mhTDP-43 antibody. Although 30 µg of protein was loaded for each sample, the band was stronger from the NEFH-TDP-43 brain due to the sample containing both the exogenous human and endogenous mouse TDP-43 proteins. The control mouse brain sample only contains the endogenous mouse TDP-43 protein. Probing the brain IP hTDP-43 samples with the mhTDP-43 antibody revealed an increasing signal with increasing amounts of the IP antibody for the NEFH-TDP-43 mice. No signal was detected in the control mouse, which is to be expected as the control mouse does not have the human TDP-43 protein present. Figure 1B shows the total protein loaded for the RIPA lysates and immunoprecipitations and shows the increasing amounts of antibody used for the IP as indicated by the IgG Heavy and Light Chains.

**Figure 1.**
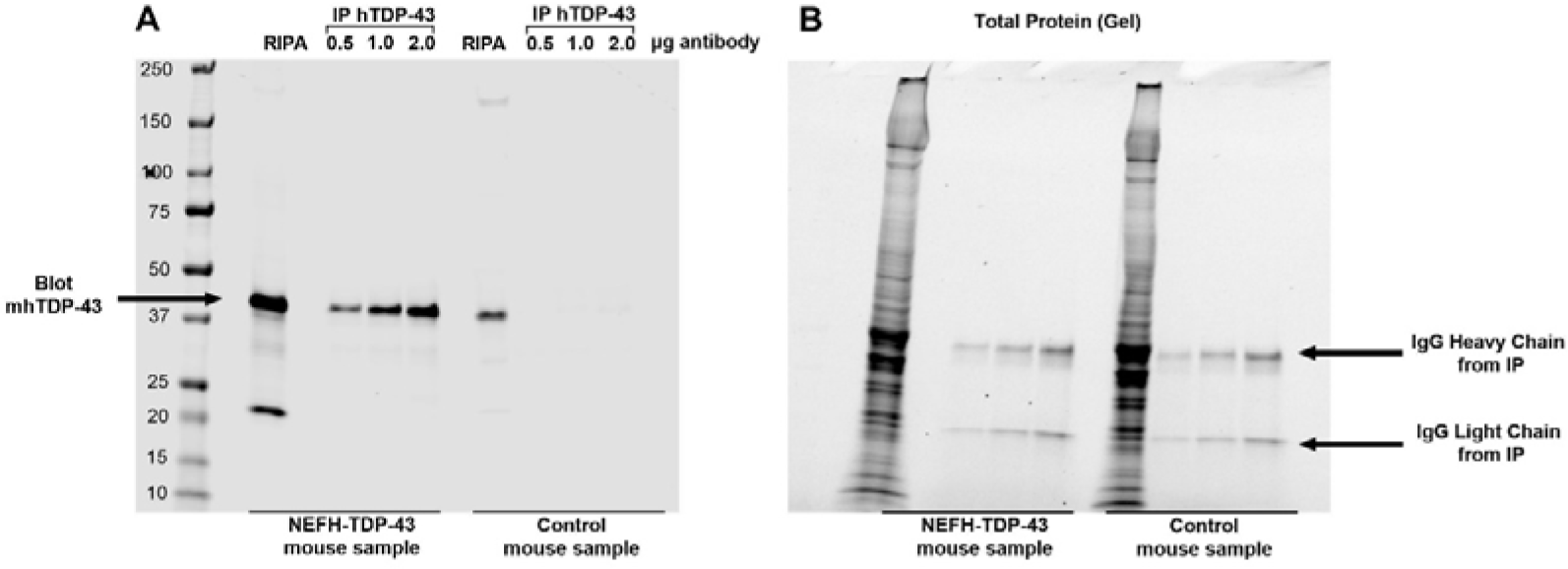
Establishing immunoprecipitation (IP) specificity using the TDP-43 antibody recognizing the human isoform (hTDP-43). (A) TDP-43 IP was performed on RIPA extracts of crushed brain tissue from the NEFH-TDP-43 and control mice using increasing amounts of the hTDP-43 antibody (0.5, 1.0 and 2.0 µg). The IP samples were then probed using a TDP-43 antibody recognizing both the mouse and the human isoforms (mhTDP-43). RIPA extracted starting material from the NEFH-TDP-43 and control mice were used as positive control samples. (B) Protein loading gel showing total protein loaded for the RIPA lysates, as well as evidence of increasing amounts of antibody used as shown by the IgG Heavy and Light Chains from the IP.

IP was performed with and without tissue crosslinking, demonstrating that crosslinking is required to maintain protein-protein interactions within our SG preparation from brain tissue (Figure 2). IP of the SG preparation was performed with a TIA-1 antibody, as TIA-1 is a known SG core protein (Kedersha et al., 2000) and binds TDP-43 (McDonald et al., 2011). The samples were then probed with the antibody recognising both the mouse and human forms of the TDP-43 protein (mhTDP-43). Figure 2A shows that after crosslinking brain samples from control and NEFH-TDP43 mice, followed by TIA1 IP, TDP-43 is detected via western blotting in the NEFH-TDP-43, but not the control sample (compare lanes 2 and 3). When cross linking is not performed prior to TIA-1 IP, TDP-43 is not detected in the NEFH-TDP-43 sample (lane 5), indicating the loss of the protein-protein interaction. Additionally, IP was performed with and without crosslinking using the antibody that recognises the human TDP-43 isoform (hTDP-43), followed by probing with the mhTDP-43 antibody. TDP-43 was detected in the NEFH-TDP-43 samples only (compare lanes 6 and 7) and was not influenced by crosslinking (compare lanes 8 and 9). This demonstrates that the crosslinking process does not disrupt the epitope for the hTDP-43 antibody.

**Figure 2.**
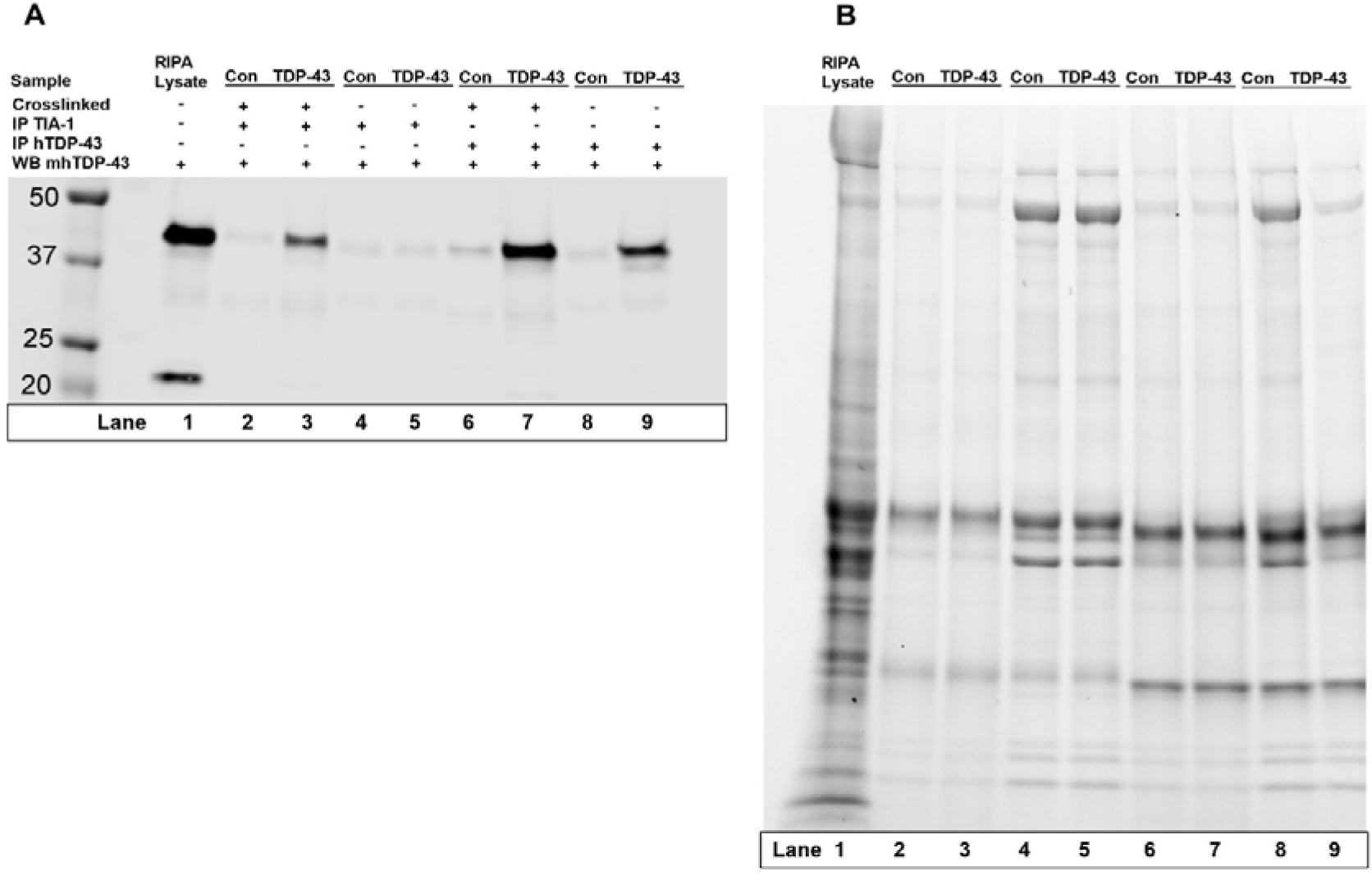
Immunoprecipitation (IP) with and without crosslinking using either TIA1 or TDP-43 antibodies. (A) IP of the SG preparation with a TIA1 antibody indicated that crosslinking is required to maintain protein-protein interactions within our SG preparation. IP of the SG preparation with the hTDP-43 antibody indicated that crosslinking does not disrupt the epitope for the hTDP-43 antibody. (B) Protein loading for all samples. Con, control mice; TDP-43, NEFH-TDP-43 mice.

### Proteomic analysis of stress granule markers

Following LC-MS analysis of the TDP-43-associated SG fractions, 4,700 quantifiable proteins were identified (Supplementary Table 1). These proteins were compared against the RNA granule database (Millar et al., 2023). Of the 4,700 quantifiable proteins 2,512 were identified as SG and RNA binding proteins (RBPs). Additionally, our proteins were compared against the mammalian SG proteome list of 540 proteins (Jain et al., 2016). From our list 383 proteins were found to be known SG marker proteins including the canonical SG initiator eIF2α, and several core SG proteins TIA-1, G3BP1, G3BP2, PABPC1 and CAPRIN1. Of the 4700 proteins, 134 proteins were significantly enriched in the brain tissue from the NEFH-TDP-43 mouse brains, when compared to the control mice (TDP-43/control = fold change >1.5, p<0.05) (Supplementary Table 2). Additionally, there were 17 proteins comparatively depleted in the SG fraction from the NEFH-TDP-43 mouse brains, when compared to the control mice (control/TDP = fold change >1.5, p<0.05) (Supplementary Table 3). Figure 3 is a representative volcano plot showing the enriched proteins in red and the depleted proteins in green.

**Figure 3.**
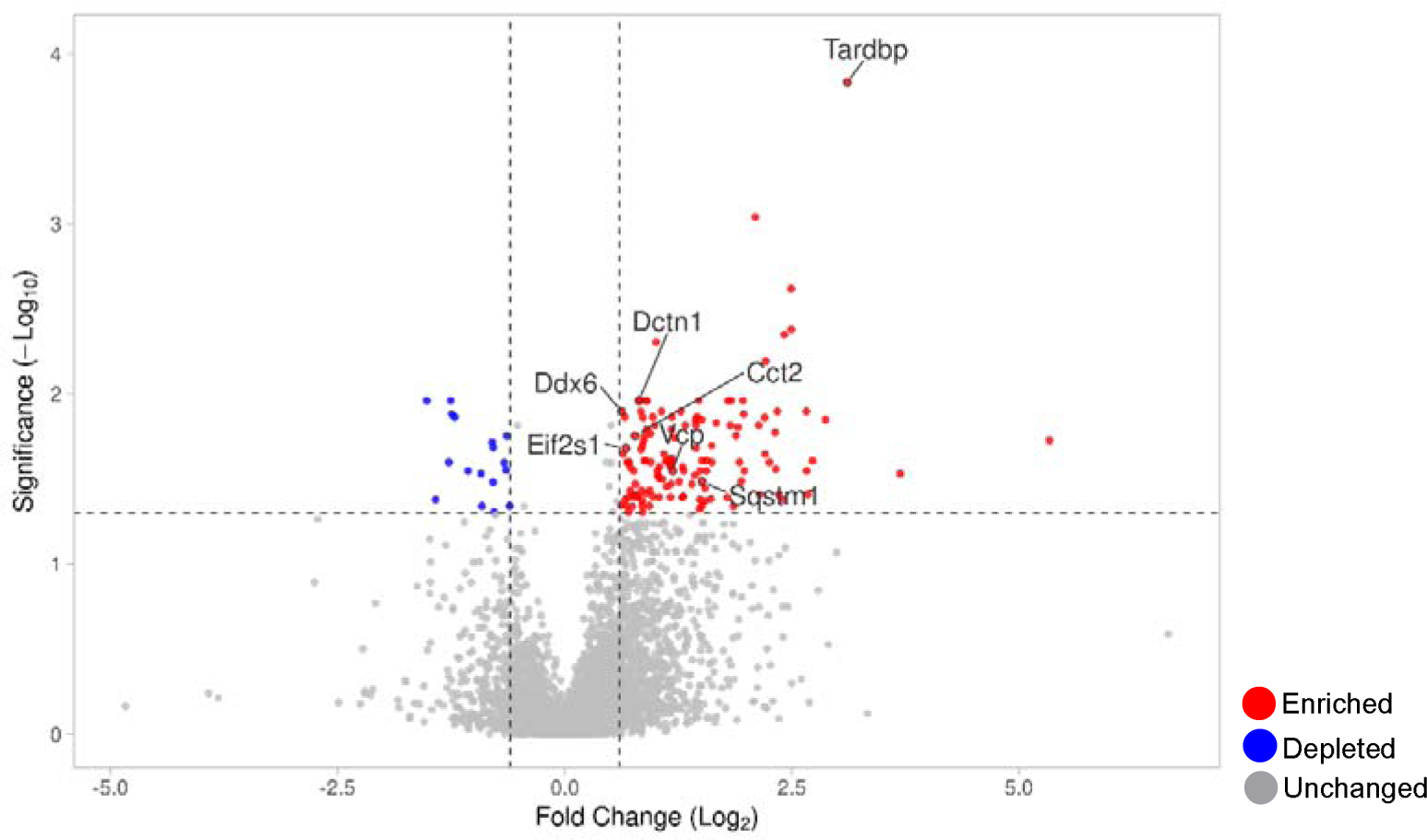
A volcano plot highlighting the enriched (red), depleted (blue) and unchanged (grey) proteins in the TDP-43-associated insoluble SG fraction isolated from the brain tissue of end-stage NEFH-TDP-43 mice. The TDP-43 protein (tardbp) is specifically highlighted to show its enrichment as are SG markers; Ddx6 (DEAD box protein 6), Dctn1 (Dynactin subunit 1), Eif2s1 (Eukaryotic translation initiation factor 2 subunit alpha), Cct2 (T-complex protein 1 subunit beta), Vcp (Valosin-containing protein) and Sqstm1 (Sequestosome-1). All proteins are labelled with the primary gene symbol for consistency.

Following comparison with the mammalian SG proteome (Jain et al., 2016), 24 of our 134 enriched proteins were known SG markers (Supplementary Table 2). Further comparison against a list of early and late stress granules markers (Hu et al., 2023) identified a further 16 SG markers, of which most were categorised as late SG markers. Therefore, 30% (41/134 proteins) of our SG preparation consisted of known SG markers (Hu et al., 2023; Jain et al., 2016). Stress granule formation is initiated by the activation of eIF2α via its phosphorylation by either PERK, HRI, GCN2 or PKR kinases (Taniuchi et al., 2016). The 134 enriched proteins and 17 depleted proteins were analysed by STRING (https://string-db.org/). The STRING analysis identified GO terms that are suggestive of activation of all four eIF2α-regulating kinases, as well as formation of SG-eIF2α complexes. Table 1 shows the GO terms, descriptors and the proteins that were identified by STRING. The GO terms are grouped by kinase/SG activation pathway. STRING was also used to visualise the protein-protein interactions between the proteins of the enriched and depleted datasets as a representation of their interactions. (Figure 4).

**Figure 4.**
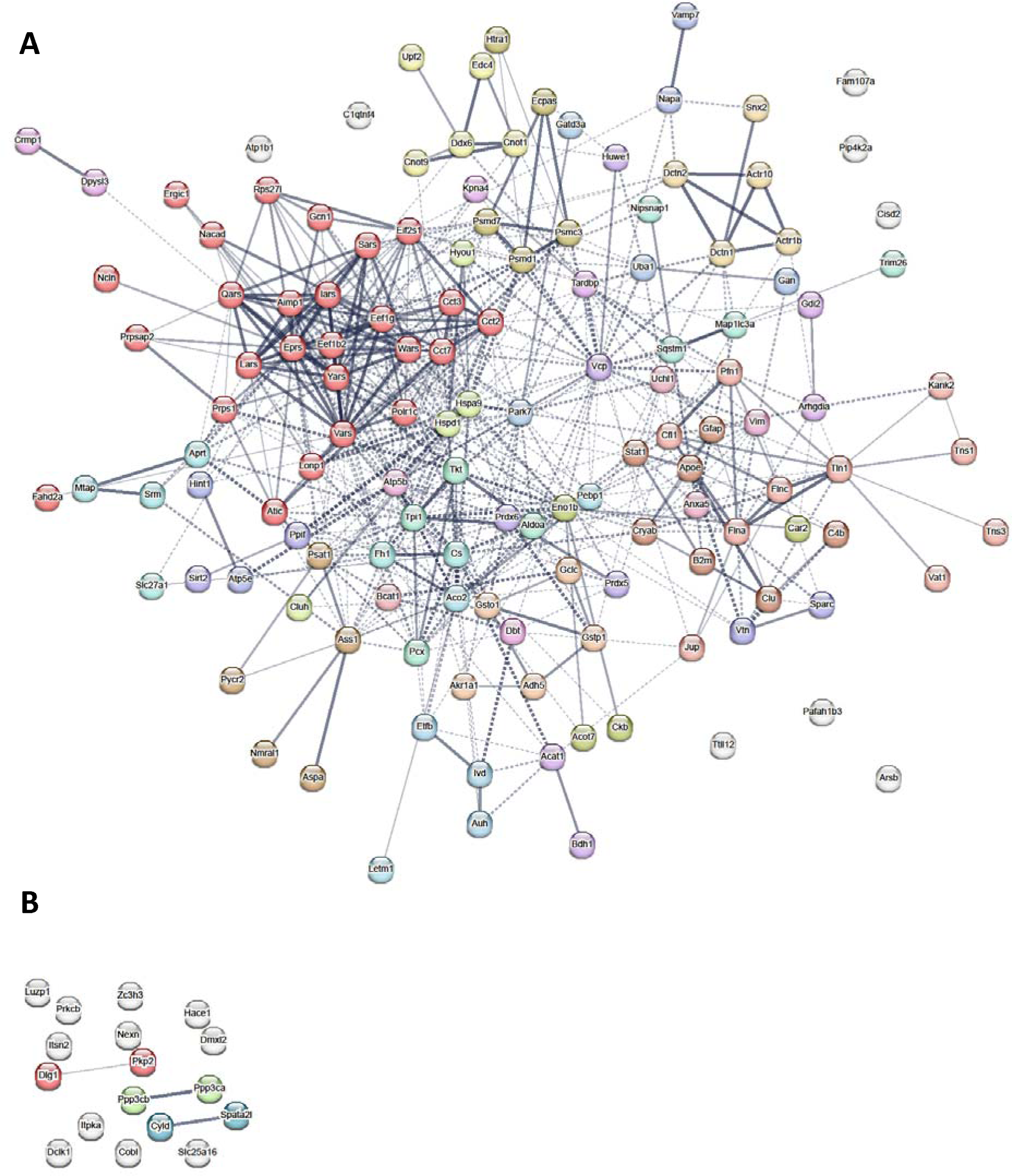
STRING representation of the interaction between the 134 enriched proteins (A) and the 17 depleted proteins (B) in the SG preparation from brain tissue of the end-stage NEFH-TDP-43 mice. The proteins are displayed with the intensity of the lines between the protein nodes representative of the confidence of the interaction. Minimum of medium confidence (>0.4). Colouring of nodes is by MCL Clustering with default settings of inflation parameter 3.

**Table 1.**
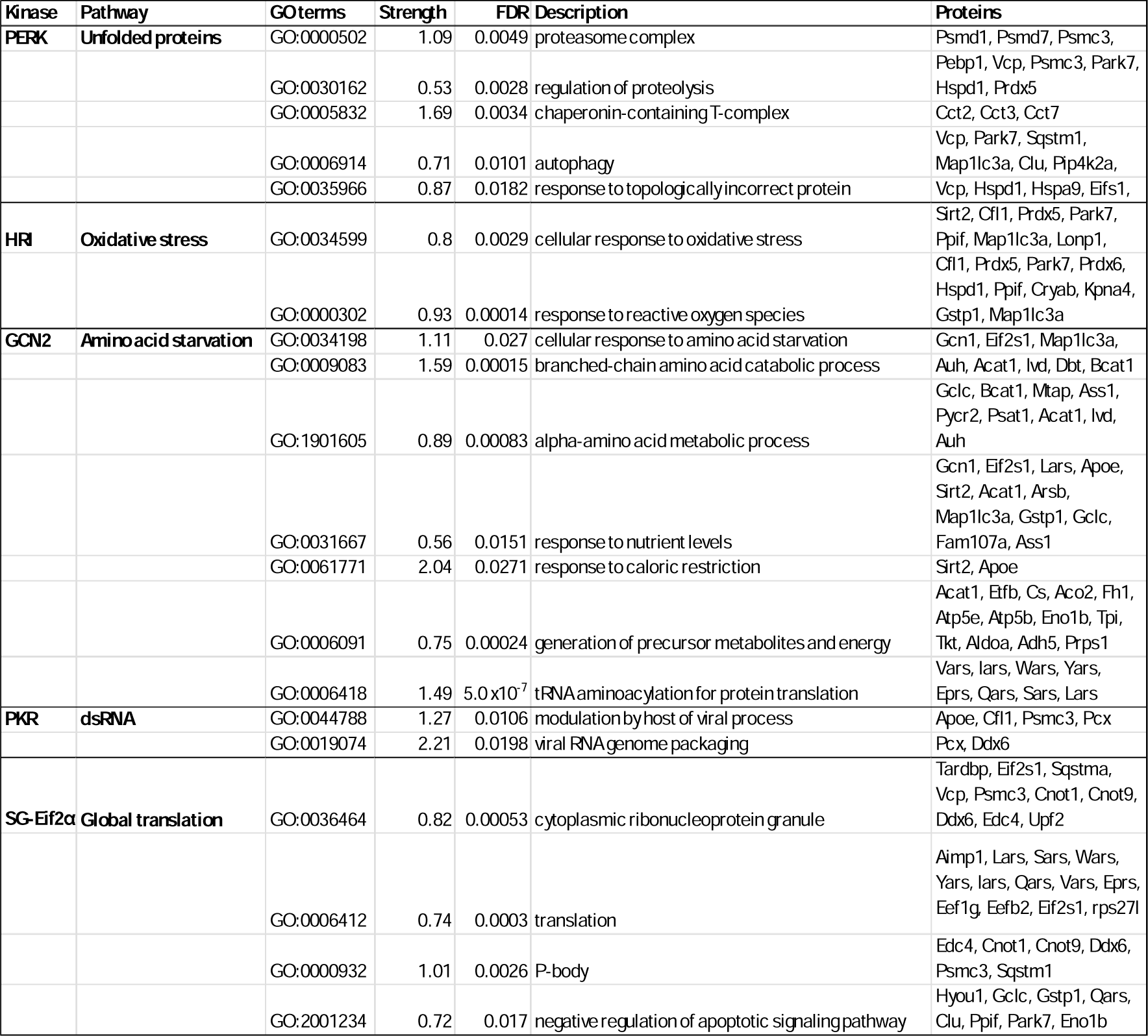
Selected enriched GO terms grouped by SG formation kinase pathways.

### Subcellular localisation of enriched proteins

Mitochondrial proteins were highlighted by the STRING analysis as the GO term “mitochondrion” (GO:0005739) included 51/134 proteins. These 51 proteins were cross checked with the mitochondrial proteome (Morgenstern et al., 2021) and the UniProt data base (https://www.uniprot.org/), indicating that 21 proteins were exclusively located to the mitochondria (Table 2). One of the proteins identified was the mRNA binding protein Clustered mitochondrial protein homologue (CLUH). It is noteworthy that 18 of the 21 mitochondrial proteins enriched were known CLUH targets (Table 2).

**Table 2.**
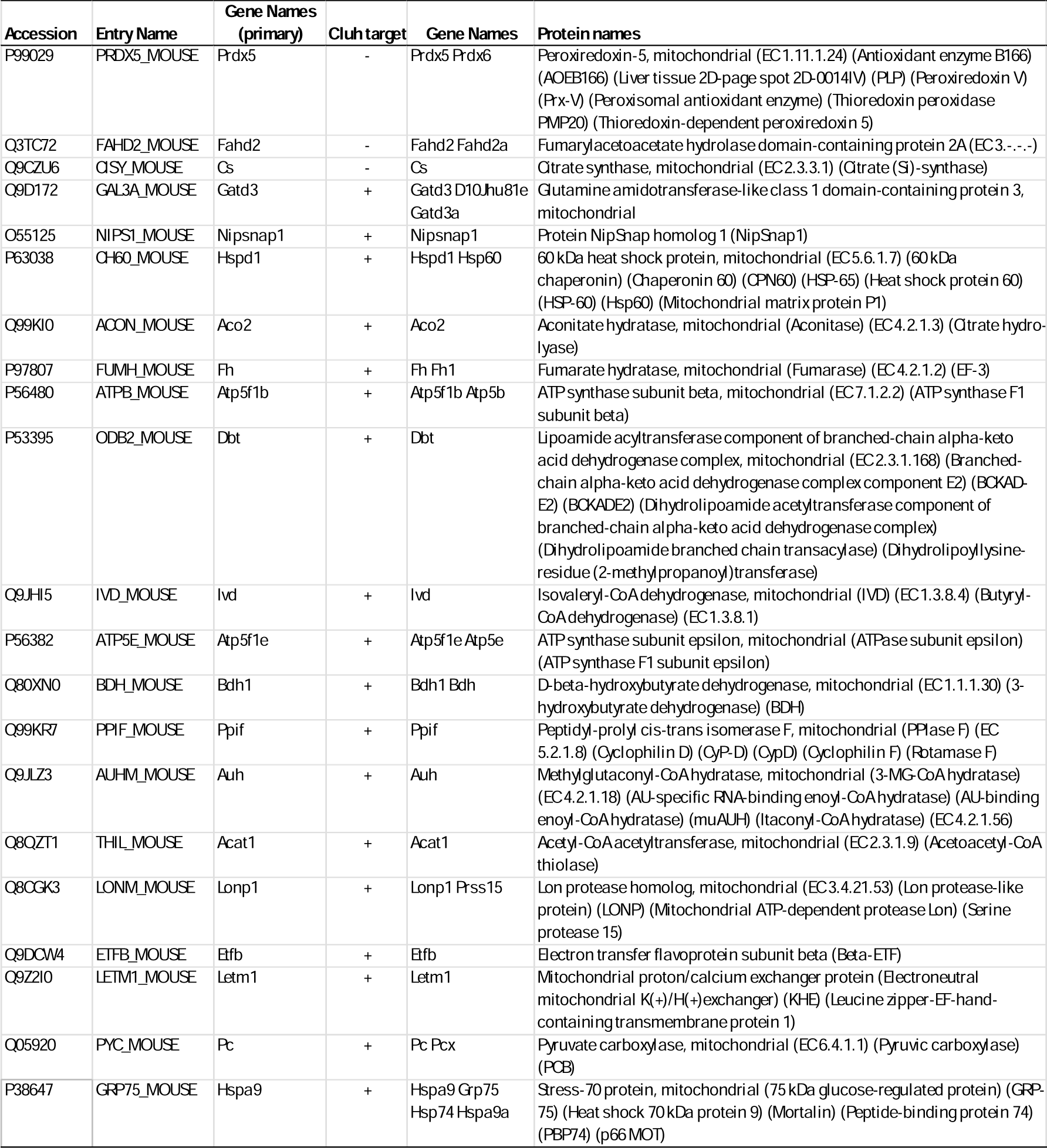
The 21 proteins exclusively located to the mitochondria.

Of the 134 proteins, it was also noted that two proteins were localised exclusively to the nucleus, (Kpna1, Polr1c) and six were exclusively secreted proteins (Apoe, B2m, C1qtnf4, C4b, Sparc, and Vtn), as based on the UniProt entry.

Figure 5 shows the interaction of these 18 CLUH targets based on STRING database analysis. Reactome pathway analysis indicated proteins related to respiratory election transport and ATP synthesis (yellow), mitochondrial import (pink), citric acid cycle (green), branched chain amino acid catabolism (purple) and utilization of ketone bodies (red).

**Figure 5.**
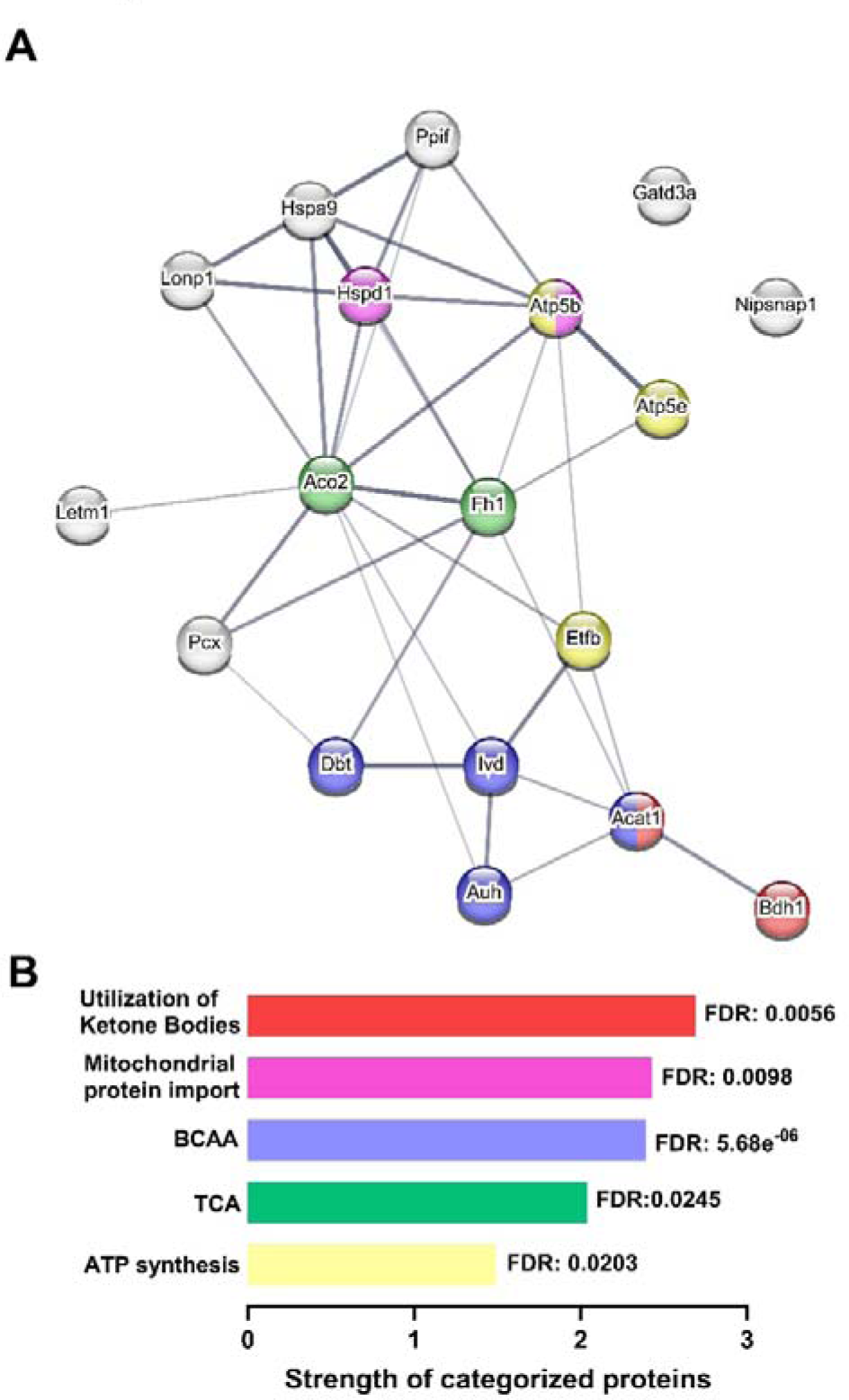
STRING representation of the interaction of CLUH target proteins. Twenty-one enriched proteins were exclusively mitochondrially located. Of these, 18 were known CLUH targets. **A.** Reactome pathway analysis indicated CLUH target proteins were enriched in pathways related to branched chain amino acid catabolism (purple), the citric acid cycle (green), respiratory election transport and ATP synthesis (yellow), the utilization of ketone bodies (red) and mitochondrial import (pink). **B.** The strength and false discovery rate of the CLUH targets Reactome pathway analysis.

Figure 6 is a summary of the results obtained from the STRING and literature analyses in relation to where many of the enriched proteins are normally located and their known biological functions.

**Figure 6:**
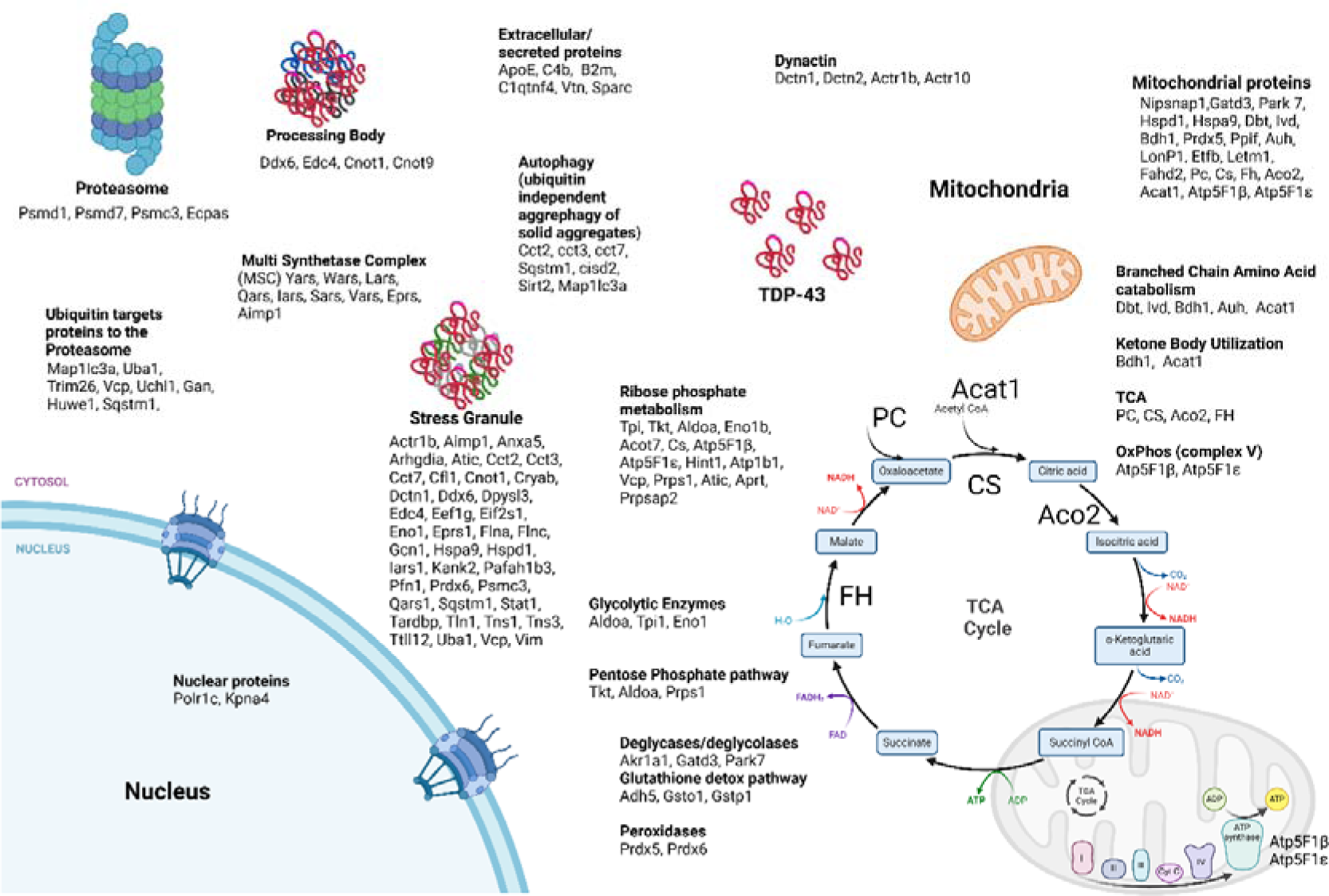
Summary of the proteins enriched in the TDP-43 immunoprecipitated stress granule preparation with their known localization and function.

### Comparison with published proteomics data derived from ALS tissue

We searched the literature to assist with placing our novel results in context of the ALS condition. Recent work by Guise *et al* (Guise et al., 2024) performed single-cell proteomic profiling of human postmortem ALS and control spinal cord MNs. Additionally, Altman *et al* (Altman et al., 2021) investigated changes in proteins from the soluble fraction isolated from sciatic nerve of NEFH-TDP-43 mice. Guise *et al* (2024) identified depletion in the soluble fraction of several pathways in the ALS MNs, including SG formation, BCAA degradation, TCA cycle, aminoacyl-tRNA synthetases (MCS), CCT and proteasome. This was of interest as these pathways were enriched in our insoluble SG fraction from end-stage NEFH-TDP-43 brain tissue (Table 1 and Figure 6). We accessed the raw dataset from Guise *et al* (Guise et al., 2024) to compare the findings from both studies. A total of 540 proteins were selected from Supplemental Table 6 from the Guise *et al* dataset that were consistently depleted in ALS, when compared to control MNs (Guise et al., 2024) (P<0.05; Fold changes > 1.5), (Supplemental Table 4). Furthermore, there were 35 identified proteins common to both datasets (Table 5). Of the 540 depleted proteins, 67 were identified as known CLUH target proteins (Supplementary Table 6).

**Table 3.**
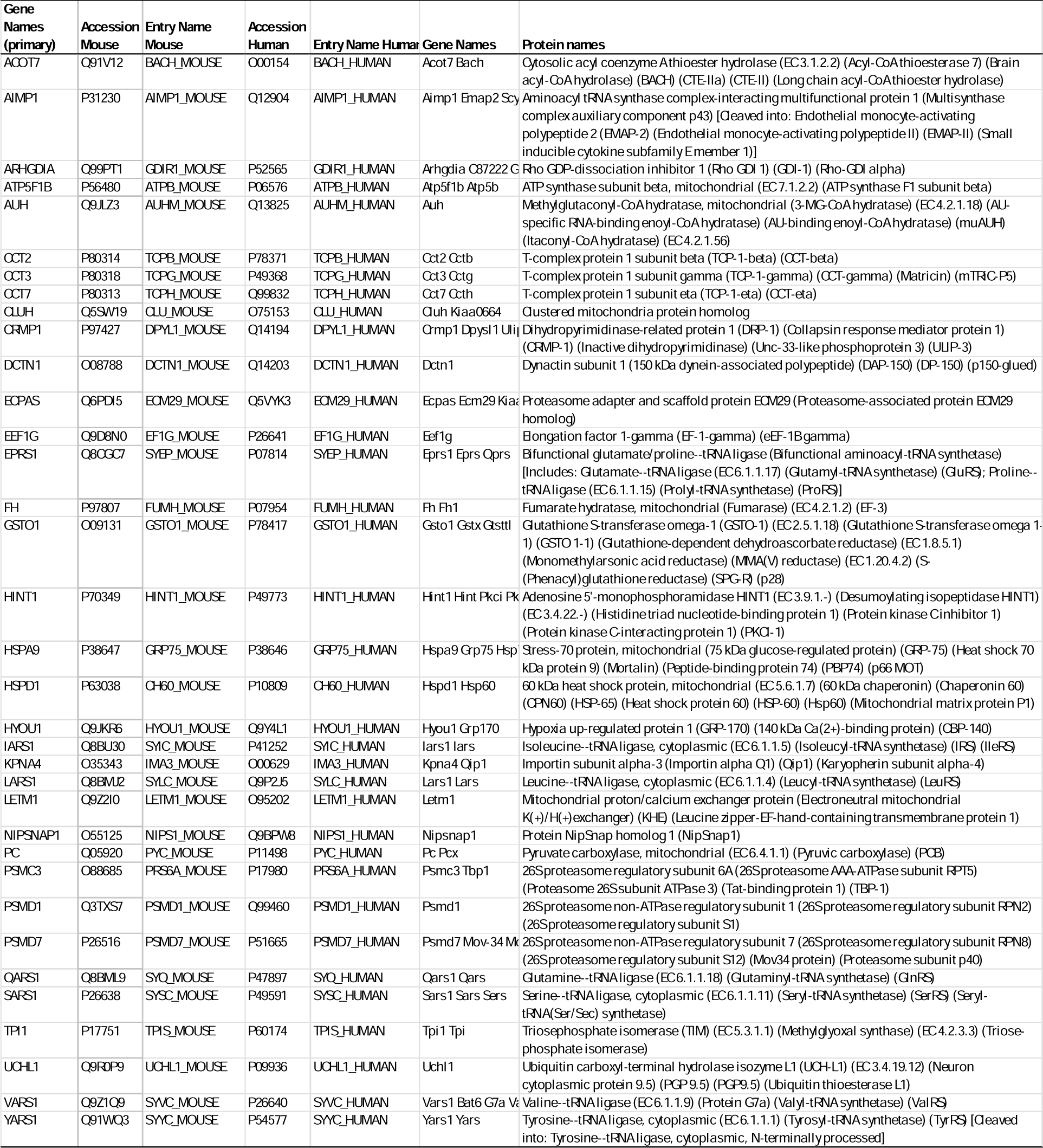
A list of the 35 proteins enriched in the insoluble SG fraction from end-stage NEFH-TDP-43 brain and depleted in the soluble protein fraction from ALS patients MNs.

The 245 depleted proteins from the soluble fraction isolated from sciatic nerve of NEFH-TDP-43 mice from Altman *et al* (Altman et al., 2021) was analysed by STRING (Supplementary Table 5). The GO term Mitochondrion covered 55% of the depleted proteins identified in the soluble fraction of the NEFH-TDP-43 sciatic nerve preparation (135 of 245) by Altman *et al* (Altman et al., 2021). Other GO terms identified from the significantly depleted proteins in the Altman *et al* dataset included Branched chain amino acid catabolism and BCAA (GO:0009083) and TCA (GO:0006099). These GO terms were also identified from the proteins enriched in our insoluble SG fraction from brain tissue. Of the 245 proteins depleted in the Altman *et al* dataset 79 are known CLUH targets (Supplementary Table 6).

Our analysis observed consistent overlap between proteins enriched in our insoluble TDP-43-associated brain SG fraction, that were also depleted in the soluble human ALS spinal MN fraction and soluble mouse sciatic nerve fractions. A Venn diagram showing the number of proteins commonly altered in the three datasets, is shown in Figure 7. When comparing the three datasets, 56 depleted targets are common between the soluble fractions from human spinal MNs and mouse sciatic nerve. Additionally, 35 proteins are common between our insoluble TDP-43-associated brain SG fraction (enriched) and the soluble human spinal MN fraction (depleted). There were four targets (PC, FH, AUH and ATP5F1B) common to all 3 datasets. Interestingly, these four are known CLUH targets.

**Figure 7.**
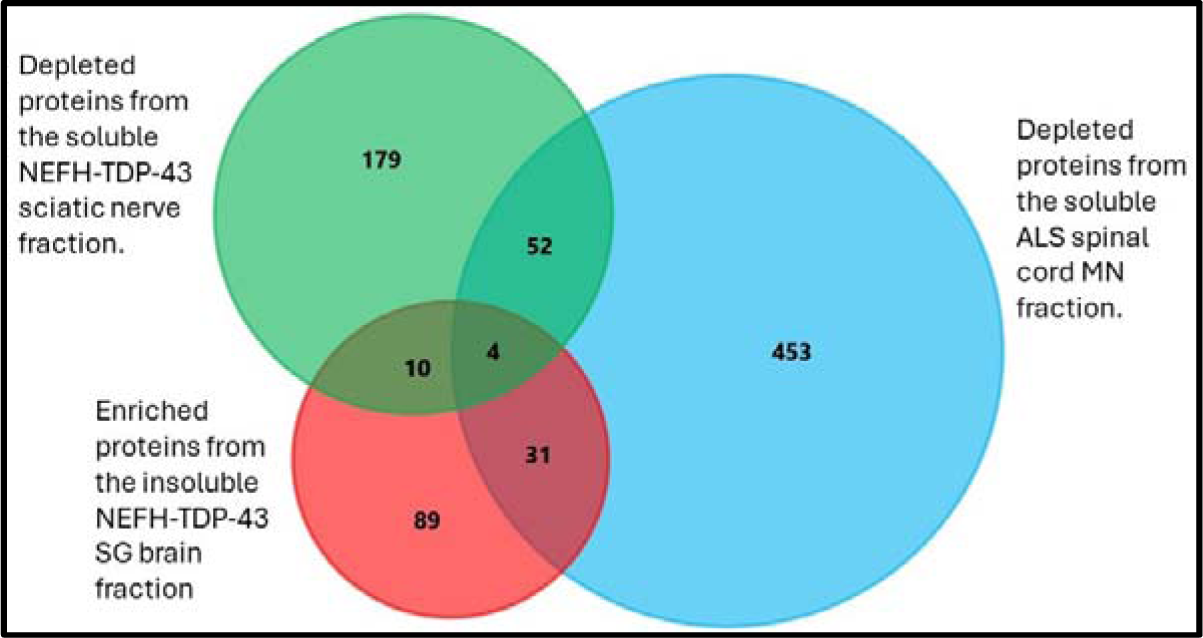
A Venn diagram showing the number of proteins that are enriched the insoluble TDP-43-associated brain SG fraction and also depleted in soluble human ALS spinal MN fraction (Guise et al. 2023) and soluble mouse sciatic nerve fraction (Altman et al. 2021).

Common overlap was also observed between the pathways regulated by CLUH and its target proteins. The common CLUH-regulated pathways enriched in our insoluble TDP-43-associated brain SG fraction, that were also depleted in soluble human ALS spinal MN fraction and soluble mouse sciatic nerve fractions are shown in Figure 8. These pathways were involved in BCCA catabolism, the citric acid cycle, respiratory electron transport/ATP synthesis and mitochondrial protein import. A table of all CLUH targets identified across all three datasets are shown in Supplementary Table 6.

**Figure 8.**
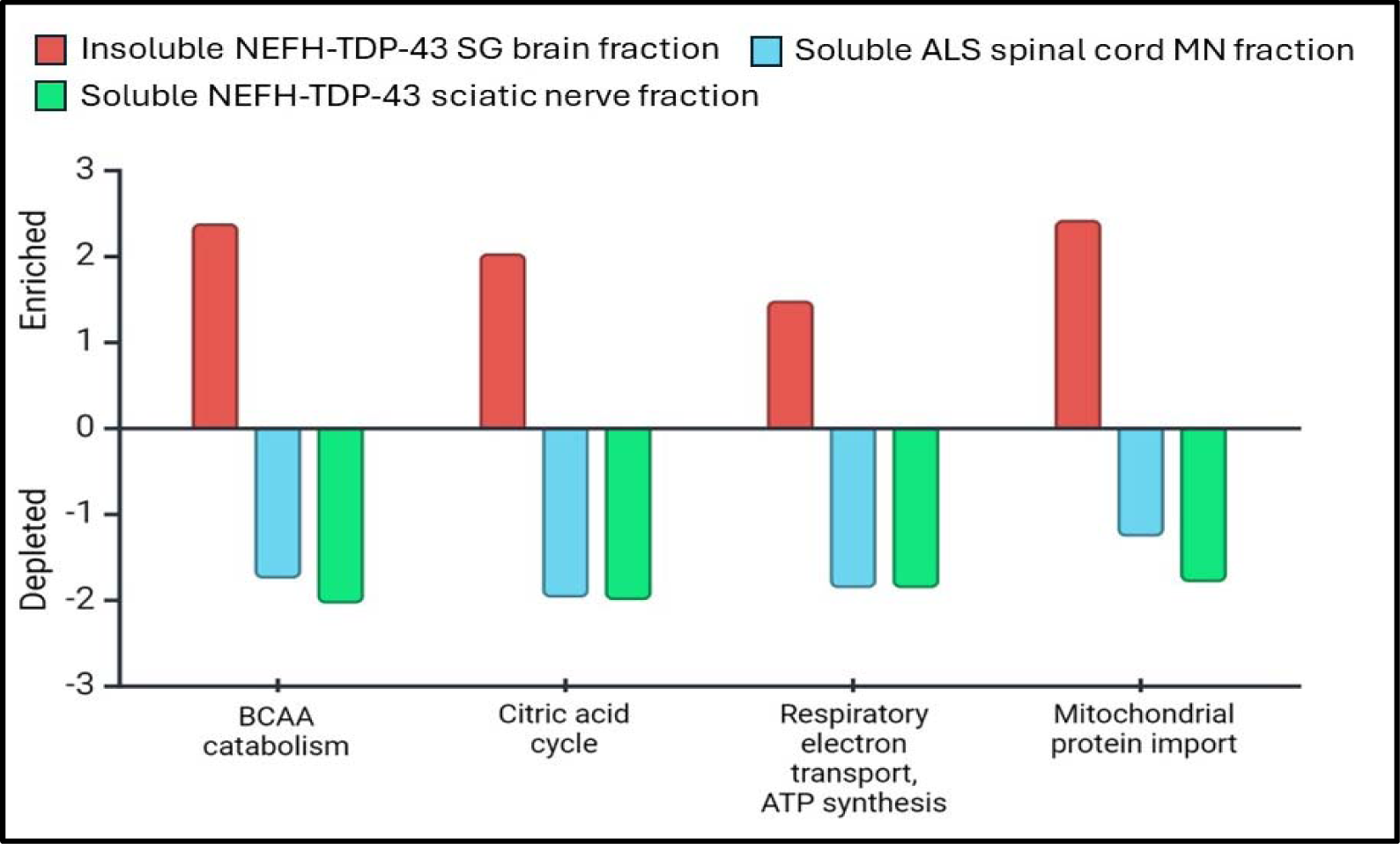
The strength and direction of the CLUH-regulated pathways identified from the proteins enriched in the insoluble TDP-43-associated brain SG fraction and depleted in soluble human ALS spinal MN fraction (Guise et al. 2023) and soluble mouse sciatic nerve fraction (Altman et al. 2021). Created with Biorender.com.

## Discussion

The aim of the work was to isolate the insoluble TDP-43 associated SGs from the brain tissue of end stage NEFH-TDP-43 mice for subsequent proteomics analysis. Almost 5000 proteins were quantified from the cross-linked SG preparations from the TDP-43 mice. Approximately half of these proteins (2512) were SG and RNA binding proteins (RBPs). This was based on comparison with the comprehensive cell culture-derived Luenfield SG database consisting of 5732 proteins (as of 25/1/2024) (Millar et al., 2023). Our *in vivo* SG dataset also contained 71% of the proteins identified by G3BP1 immunoprecipitation from formaldehyde cross-linked, sodium arsenite stressed Human U-S O2 cells (Jain et al., 2016). This supports the specificity of our modified formaldehyde cross-linking and TDP-43 (a core SG protein) immunoprecipitation protocol for use in brain tissue, that had previously been used in cultured cells (Jain et al., 2016). Therefore, to the best of our knowledge, this is the first time an insoluble crosslinked TDP-43-associated SG fraction has been isolated from brain tissue of an ALS mouse model and its proteome reported.

The data presented here provides an end-stage snapshot of insoluble cytoplasmic TDP-43-associated SGs and other RBP partners. Analysis of the enriched proteins and an understanding of their respective pathways, and possible interactions, will shed light on potential ALS disease modifying mechanisms. From the ∼5000 proteins quantified, a subset of 134 proteins were enriched and 17 proteins depleted, in the end-stage TDP-43 mouse brain tissue, compared with the control mice. Of these enriched proteins, 41 proteins were SG markers, more than a third of which were late SG markers (Hu et al., 2023; Jain et al., 2016). Late stress granule markers associated with the proteasome, processing bodies, multi-synthase complex (MCS), and the profilin 1 complex were of particular interest as it suggests sustained SG recruitment (Hu et al., 2023).

Stress granule formation is initiated by phosphorylation of eIF2α by either PERK, HRI, GCN2 or PKR depending on the stressor (Taniuchi et al., 2016). In the NEFH-TDP-43 mice, over-expression of cytoplasmic mutated hTDP-43 protein is expected to activate SG formation via the unfolded protein response (UPR) detection of ER stress-activation and subsequent activation of the PERK pathway (Walker et al., 2013). This is supported by the enrichment of GO terms related to unfolded proteins (See table 1). Mitochondrial import of TDP-43 increases mitochondrial ROS production (Wang et al., 2019). Therefore, it was not unexpected to see enrichment of proteins associated with oxidative stress and ROS. This is consistent with activation of SGs via the HRI pathway (Basu et al., 2017). Enriched proteins associated with GO terms for amino acid starvation and nutrient levels supports SG activation via the GCN2 pathway (Hinnebusch & Fink, 1983). Additionally, there was an enrichment of GCN1, a direct activator of GCN2 (Cambiaghi et al., 2014), which detects the collision of ribosomes that are stalled during translation when amino acids become deficient (Lokdarshi & von Arnim, 2022). We also observed the enrichment of eight members of the Multi Synthetase complex (MCS), which act as an amino acid sensor to provide individual amino acids during protein translation (Hyeon et al., 2019), further suggesting GCN2 activation. Another novel observation was the enrichment of Polr1c in the TDP-43-associated SG fraction. The role of Polr1c is usually in the nucleus, however it also functions as a cytoplasmic DNA sensor. It activates the PKR-SG pathway, typically in response to viral infection (Chiu et al., 2009; Kessler & Maraia, 2021; Matsumiya et al., 2023). In the NEFH-TDP-43 mouse model, we suggest that SG formation may occur in response to TDP-43 entry into the mitochondria (Altman et al., 2021) and subsequent mitochondrial DNA release into the cytoplasm through the mitochondrial Permeability Transition Pore (mPTP) (Yu et al., 2020). The enrichment of Cyclophilin D (PPIF), which is part of mPTP (Yu et al., 2020), in the TDP-43-associated SG fraction supports this. Cytoplasmic Polr1c could convert the mitochondrial DNA to dsRNA and therefore activate PRK. It was unexpected to see evidence of activation of all four of the Eif2α stress granule activation pathways and this raises the possibility of a stress induced feedback loop leading to sustained SG formation.

The presence of KPNA4 (also known as Importin α3) in our insoluble SG fraction is also unexpected as it acts as the nuclear import receptor for TDP-43. The mutations introduced into the nuclear localisation sequence (NLS) of TDP-43 abolish binding to Importin α3 (Park et al., 2020). This points to a couple of possibilities, including a potential limitation of formaldehyde cross-linking. Formaldehyde requires direct contact between molecules for a bond to be formed. However, cross-linking can also occur between molecules within a complex (Klockenbusch et al., 2012). This means that proteins enriched by immunoprecipitation with TDP-43 may not be in direct contact with TDP-43 but may be part of a complex with TDP-43. With this limitation in mind, the possibility that KPNA4 was recruited to the SG complex by another SG protein needs to be considered. For this to be mechanistically important, it would need to be early in the SG formation pathway. Indeed, KPNA4 can interact with PKR and eIF2α (Piazzi et al., 2019), the latter enriched in our SG dataset and a central regulator of SG formation (Kedersha et al., 2005). Recruitment of KPNA4 into the SG fraction from the transgenic NEFH-TDP-43 mice is most likely independent of TDP-43 due to its mutated NLS within the induced human TDP-43 protein. This observation is important and suggests a possible mechanism for TDP-43 cytoplasmic accumulation under pathophysiological ALS conditions. Cytoplasmic accumulation of TDP-43 is considered a hall mark of ALS and the mechanism by which this occurs remains unknown. However, there is evidence that disruption of the interaction of TDP-43 with its import receptors is a key part of this mechanism and SGs appear to be an integral part of this process (Prpar Mihevc et al., 2017; Zhang et al., 2018). In fact, in familial C9orf72 ALS, PKR is activated by RNA products from the random insertions in the C9orf72 gene (Parameswaran et al., 2023). This phosphorylates eIf2α and recruits Importin α3 (KPNA4) into SGs, resulting in TDP-43 cytoplasmic accumulation (Khosravi et al., 2017). Additionally, PKR inhibitors prevent eIf2α phosphorylation in C9orf72 and attenuate disease progression (Zu et al., 2020). As such, the PKR inhibitor Metformin, in addition to being used for the treatment of type II diabetes, is currently in clinical trial for fALS C9orf72 (Zu et al., 2020).

An unexpected and novel finding was the enrichment of the RNA binding protein CLUH, and a group of 18 CLUH targets (Gao et al., 2014; Schatton et al., 2017). To the best of our knowledge, CLUH regulation has not been highlighted previously in the ALS field. However, the presence of CLUH in our end-stage SG preparation may have considerable relevance for ALS pathogenesis, specifically in relation to mitochondrial function, MN and NMJ degeneration and metabolic flexibility. CLUH is the mRNA binding protein responsible for mitochondrial transport of nuclear encoded mitochondrial mRNA (Gao et al., 2014; Schatton et al., 2017; Wakim et al., 2017). CLUH is also required for neuronal development and motor neuron integrity (Zaninello et al., 2023). In response to glucose and amino acid starvation-induced stress, CLUH and its bound mRNAs form large RNP granules (Pla-Martin et al., 2020; Yang et al., 2022). These CLUH granules are formed as a survival mechanism to protect mRNAs from degradation and attenuate cellular anabolism (Pla-Martin et al., 2020; Yang et al., 2022). The loss of CLUH function alters the mitochondrial proteome, impairs mitochondrial morphology and respiration, and elevates ROS production (Gao et al, 2014, Cox, Spradling 2009; Schatton Pla-Martoin, 2017; Sen, Cox, 2022; Wakim et al., 2017). Cultured liver cells from CLUH KO mice are highly glucose dependent and fail to shift from glucose to fat metabolism under starvation conditions (Schatton et al., 2017; Wakim et al., 2017). This highlights a role of CLUH in maintaining metabolic flexibility.

The CLUH targets identified in the TDP-43-associated SG fraction were enriched for the catabolic enzymes Acat1, Bcat1, Auh, Dbt, Ivd and Bdh1. These enzymes belong to the branched chain amino acid and ketone body pathways (Hwang et al., 2022; Jiang et al., 2022; Neinast et al., 2019). This suggests recruitment of CLUH to the TDP-43-associated SG fraction is a metabolic response to starvation in the end-stage brain of the NEFH-TDP-43 mice. The brain is usually protected from starvation and utilises glucose as a substrate (Harris et al., 2012; Oyarzabal & Marin-Valencia, 2019). However, it can use branched chain amino acids and ketone bodies, specifically leucine and β-hydroxybutyrate respectively (Divakaruni et al., 2017; Jensen et al., 2020). Although speculative, CLUH recruitment into the TDP-43-associated SG fraction would diminish its physiological role and contribute to the excessive impairment in mitochondrial function, as well as axonal and NMJ degradation seen in ALS (Altman et al., 2021; Zaninello et al., 2023). Identifying the point during ALS disease progression when CLUH is recruited to TDP-43-associated granules will be important to understand mechanisms of disease progression and potentially provide an opportunity for therapeutic intervention.

Mitochondrial dysfunction and impaired ATP production is well established in ALS and a consequence of multiple cellular disturbances (Muyderman & Chen, 2014; Zhao et al., 2022). Disrupted oxidative phosphorylation activates the pentose phosphate pathway (PPP) to generate cytoplasmic ATP (Skinner et al., 2023). Cell starvation, as indicated by CLUH recruitment into TDP-43-associated granules, would attenuate the provision of glycolytic and PPP substrates required for ATP production. Under such conditions ribose salvaging from RNA is activated as a carbon source for ATP generation. Our SG protein dataset was enriched for ribose salvaging from RNA for cytoplasmic ATP generation (GO:0019693 Ribose phosphate metabolic process). Furthermore, enrichment of Prps1, Tkt and Aldoa in our SG fraction suggests an impact on the PPP. Considering the fundamental role of SG formation, these PPP enzymes would be recruited into SGs to protect the untranslated mRNA. We speculate that the SG response and the CLUH granule response would prevent SG resolution, as this is an ATP dependent process (Jain et al., 2016).

This raises the possibility that severe ALS pathology could be explained in terms of stress-related feedback loops resulting in persistent and progressive SG and CLUH granule recruitment with mitochondrial disruption leading to neuronal starvation and ATP crisis. Figure 9 presents a hypothetical mechanistic framework, developed from our results and other published literature, linking acute TDP-43 cytoplasmic aggregation and SG formation to mitochondrial damage, sustained eIF2α activation and chronic SG formation, to eventual CLUH granule recruitment and MN and NMJ deterioration.

**Figure 9.**
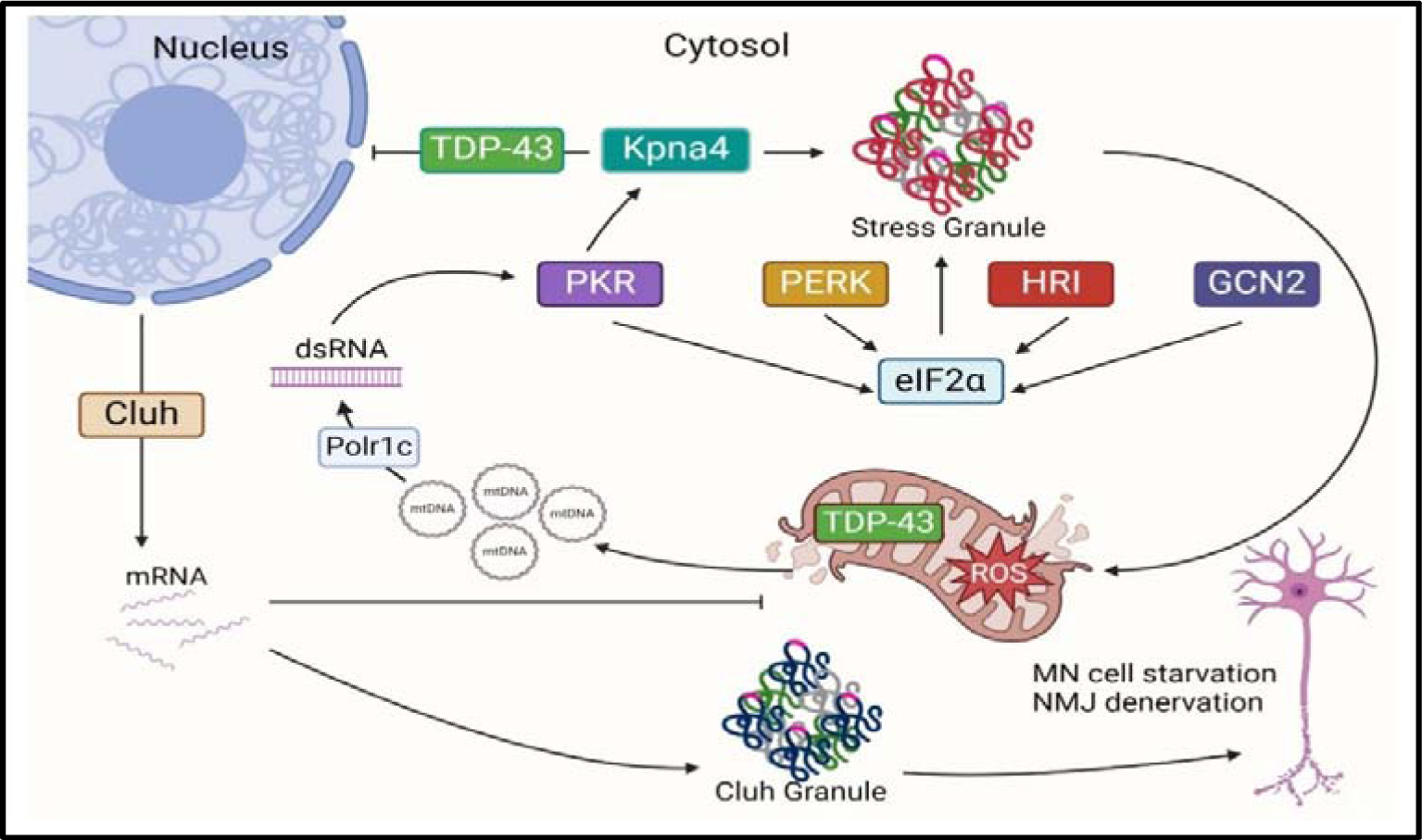
Theoretical model depicting the relationship between cytoplasmic TDP-43 and aggregation, SG formation, mitochondrial impairment and eventual NMJ and MN damage. TDP-43 accumulation and aggregation activate SG formation via multiple kinases upstream (PKR, PERK, HRI and GCN2) of eIF2α. The accumulative activation of these multiple kinase pathways and sustained SG formation would increase disease progression and severity due to persistent mitochondrial damage and mitochondrial DNA release into the cytoplasm, forming an escalating SG response. From the initial activation of the unfolded protein response via PERK, eIF2α phosphorylation activates the integrated stress response. This would cause a global halt to translation and suppression of apoptosis as the cells attempt to adapt to the stress. This recruits KPNA4 (Importin α3) into SGs, leading to the cytoplasmic accumulation of TDP-43. TDP-43 mitochondrial import would occur as a potential mechanism to remove excess TDP-43 via the ATP-dependent degradative action of LonP1, or to regulate mitochondrial metabolism. Excess mitochondrial TDP-43 would contribute to mitochondrial dysfunction, accumulation of ROS and activation of the HRI SG pathway. Prolonged cellular stress and eventual metabolic inflexibility would sequester CLUH (and its target mRNA/proteins) to the SG fraction to attenuate cellular anabolism. This would block mitophagy via loss of the CLUH target nipsnap1, causing damaged mitochondria to accumulate, followed by cytoplasmic mitochondrial DNA accumulation via the mPTP, as well as NMJ denervation and MN cell starvation. The cytoplasmic DNA is converted to RNA by Polr1c, which activates PKR and escalating the SG response. Created with Biorender.com.

The positive effect of uridine supplementation on translation in CLUH KO motoneurons (Zaninello et al., 2023), supports the notion of stress-related feedback loops leading to neuronal starvation and ATP crisis. From a therapeutic perspective, uridine can substitute for ribose in the PPP, potentially sparing the RNA (Skinner et al., 2023). Uridine prevents ATP loss and *in vitro* cell death in glucose starved neurons in a dose dependent manner (Choi et al., 2008). In the context of ALS, uridine treatment extends the survival of SOD1G93A ALS mice and attenuates body weight loss, also in a dose dependent manner (Amante et al., 2010). This supports the notion of neuronal starvation in ALS. Oral uridine is poorly absorbed. However, the uridine pro-drug triacetyluridine, which is used for the treatment of 5-FU toxicity, results in supraphysiological uridine levels and is well absorbed by the gastrointestinal tract (Garcia et al., 2005). Triacetyluridine may therefore have a neuroprotective affect in ALS-induced neuronal starvation.

Analysis of the proteins depleted from the TDP-43-associated SG fraction identified several proteins with complimentary functions to the enriched protein data set, strengthening our interpretations. These depleted proteins included Dclk1, Zc3h3, PPP3CA and CLYD. Dclk1 is a kinase involved in the enhancement and inhibition of anterograde and retrograde transport, respectively in axons (Nawabi et al., 2015). Zc3h3 is a zinc finger protein required for nuclear export of mRNA and regulation of mRNA decay (Silla et al., 2020). Of significant interest, and in alignment with the enrichment of the CLUH-related granules, was the depletion of the calmodulin-dependent calcineurin A phosphatase (PPP3CA). PPP3CA dephosphorylates DNM1L (DRP1 was detected, but not enriched in our data set) in response to increased Ca2+ from mitochondrial depolarization (Cereghetti et al., 2008). Dephosphorylated Drp1 translocates from the cytoplasm to the mitochondria, via a CLUH-dependent mechanism (Yang et al., 2022). Here it triggers mitochondrial fission to separate damaged mitochondria, potentially in preparation for mitophagy (Yang et al., 2022; Zerihun et al., 2023). There was depletion in the Ubiquitin carboxyl-terminal hydrolase (CLYD) protein. CLYD promotes microtubule stabilisation by inhibiting HDAC6 deacetylation of alpha-tubulin (Wickstrom et al., 2010). Interestingly, there was enrichment for Tubulin deacetylation (GO:0090042), including the Tubulin deacetylase, Sirt1. Combined, these observations are consistent with the importance of tubulin acetylation in axonal transport in ALS (Guo et al., 2017).

Analysis of the soluble ALS patient-derived MN dataset (Guise et al., 2024) and comparison with our results, sheds some light on how TDP-43 associated SGs may have a broad impact on multiple cellular pathways. Several functional networks enriched in our insoluble TDP-43 associated SGs brain dataset were downregulated in the soluble spinal cord MN dataset (Guise et al., 2024), including proteasome complex, chaperonin-containing T-complex, tRNA aminoacylation for protein translation, TCA and oxidative phosphorylation. These inverse changes in the functional networks from insoluble brain and soluble MN samples is consistent with the role of RBPs and RNA granules in the transport of mRNA for local translation in distal axons of MNs (Altman et al., 2021; Piol et al., 2023). For example, CLUH has a specific role in the transport and stability of nuclear encoded mitochondrial mRNA in MNs and maintenance of the NMJ (Gao et al., 2014; Schatton et al., 2017; Zaninello et al., 2023). The recruitment of CLUH and CLUH target mRNA to the insoluble TDP-43-associated SG fraction in the brain may result in depletion of these same molecules in the functional soluble component in distal axons of MNs. Indeed, observations made *ex vivo,* that CLUH and 64 CLUH target proteins were depleted in the soluble fraction of ALS patient MNs is striking (Guise et al., 2024). Importantly, five of the depleted CLUH target proteins (Gao et al., 2014; Schatton et al., 2017) were also our enriched dataset (Nipsnap1, Hspd1, Atp5f1b, Letm1 and Pc).

Observations made in the soluble ALS patient-derived MN dataset (Guise et al., 2024) also support this, including downregulation of Reactome pathways and GO terms relating to cellular response to starvation, response of EIF2AK4 (GCN2) to amino acid deficiency, response to ER stress (PERK), as well as downregulation of both PKR (E2AK2) and KPNA4 and enzymes involved in branched chain amino acid catabolism.

Reanalysis of the Altman *et al* dataset of depleted proteins in the soluble fraction from sciatic nerves of the NEFH-TDP-43 mice, which is heavily skewed for mitochondrial proteins, revealed that 33% of the total depleted dataset are known CLUH targets. These two axon-derived depleted protein datasets support a link between the formation of insoluble SG aggregates and protein depletion in the soluble protein pool. Furthermore, the datasets highlight a likely role of CLUH in this depletion. STRING analysis identified retrograde dynactin motor components enriched in our insoluble SG dataset (Dcnt1, Dctn2, Actr1b and Actr10) and depleted in the soluble ALS patient MNs samples (Dctn1, Actr1a, Dync1li2, Dync1i1, Dynll1, Dync1h1, and Dync1li1). Additionally, imaging experiments of TDP-43-containing granules conducted in ALS patient iPSC-derived neurons, shows a specific increase in retrograde axonal transport (Alami et al., 2014). This suggests the mechanism of depletion of CLUH target proteins from MNs by TDP-43-associated granules is via enhanced retrograde transport.

Here we have observed that the insoluble TDP-43-associated SG proteome from end-stage NEFH-TDP-43 brain tissue is enriched with not only well-studied SG markers, but also enriched with numerous other proteins. Integration of these unexpected SG-bound proteins suggests activation of multiple SG forming pathways, mitochondrial damage and cytoplasmic presence of mitochondrial DNA; the later forming a loop for sustained SG formation. We have provided a hypothetical mechanistic framework, based on the interpretation of our results and in conjunction with published literature, demonstrating how alterations in the pathways may intersect to create a toxic environment resulting in MN and NMJ degeneration (Figure 9). The presence of CLUH is of significant interest considering its role in mitochondrial and neuronal homeostasis and should be considered as a novel target regulating ALS pathogenesis. Understanding the temporal formation and composition of this SG fraction will further identify key disease modifying proteins and pathways influencing ALS.

## Acknowledgements

This work was supported by funding provided by FightMND (IP-158) and Institute for Physical Activity and Nutrition, School of Exercise and Nutrition Sciences.

